# Obesity differentially effects the somatosensory cortex and striatum of TgF344-AD rats

**DOI:** 10.1101/2024.01.22.576454

**Authors:** Minhal Ahmed, Aaron Y. Lai, Mary E. Hill, Jessica A. Ribeiro, Ashley Amiraslani, JoAnne McLaurin

## Abstract

Lifestyle choices leading to obesity, hypertension and diabetes in mid-life contribute directly to the risk of late-life Alzheimer’s disease (AD). However, in late-life or in late-stage AD conditions, obesity reduces the risk of AD and disease progression. To examine the mechanisms underlying this paradox, TgF344-AD rats were fed a varied high-carbohydrate, high-fat (HCHF) diet to induce obesity from nine months of age representing early stages of AD to twelve months of age in which rats exhibit the full spectrum of AD symptomology. We hypothesized regions primarily composed of gray matter, such as the somatosensory cortex (SSC), would be differentially affected compared to regions primarily composed of white matter, such as the striatum. We found increased myelin and oligodendrocytes in the somatosensory cortex of rats fed the HCHF diet with an absence of neuronal loss. We observed decreased inflammation in the somatosensory cortex despite increased AD pathology. Compared to the somatosensory cortex, the striatum had fewer changes. Overall, our results suggest that the interaction between diet and AD progression affects myelination in a brain region specific manner such that regions with a lower density of white matter are preferentially effected. Our results offer a possible mechanistic explanation for the obesity paradox.

## 2 Introduction

Alzheimer’s disease (AD) is the most common form of dementia with over 50 million patients internationally in 2019^1^. In the last 20 years, the contribution of lifestyle factors on dementia risk has been increasingly reported with the impact of obesity on AD development and progression recognized^2,3^. With the obesity epidemic underway, it is important to understand the link between obesity, age and AD^4^. In humans, obesity and a sedentary lifestyle contributes to neurocognitive decline^5^. Numerous studies in patients have also linked mid-life obesity to increased AD risk and cognitive decline^6–9^. In contrast, studies have found that obesity in late-life reduces AD risk and protects against neurocognitive decline as AD progresses thus decreasing the severity of disease in patients^10–13^. This dichotomy is known as the obesity paradox and remains poorly understood.

Genetic and dietary models have been used to examine obesity in animals with conflicting results^14,15^. In relation to AD, some studies have found that diet-induced obesity accelerates AD in rodents^16,17^, however, other studies have found that diet-induced obesity has no impact on AD or can even protect against AD in rodent models^18,19^. These combined studies fed rodents with high fat and/or high carbohydrate diets from a young age^17–20^. Here, we model obesity utilizing a varied high-carbohydrate, high-fat (HCHF) diet in which food items are rotated to keep rats engaged with the diet as opposed to standard chow (CHOW)^20^. A varied HCHF diet has been found to cause obesity in rat models after a 12-week feeding period^20^. We utilized the TgF344-AD (TgAD) rat model which overexpresses human amyloid precursor protein containing the Swedish familial mutation and presenilin 1 ΔE9 mutation under the prion promoter^21^. At nine months of age, TgAD rats begin to exhibit cognitive decline, amyloid-,B (A,B) accumulation, and vascular dysfunction representing an early stage of AD^22^. At twelve months of age, TgAD rats have more extensive cognitive decline, increased A,B plaques, cerebral amyloid angiopathy, and exhibit significant tau inclusions representing established disease^21^. Rats were fed a HCHF diet beginning at nine months of age up until twelve months of age to examine the interaction between HCHF and the presence of AD symptomology.

We hypothesized that different brain regions would be differentially effected by HFHC diet. The two brain regions chosen were the striatum and the somatosensory cortex (SSC), both of which exhibit decline of function in AD^23,24^. The striatum is involved in learning and planning, and is primarily composed of white matter^25^. The somatosensory cortex is involved in motor and haptic memory, and is primarily composed of gray matter^26^. By choosing these two regions, we examined whether a HCHF diet had differential impacts on white and gray matter. To this end, we conducted pathological assays to examine the interactive effect of a HCHF diet and AD progression on the striatum and the somatosensory cortex.

## 3 Results

Sex differences were not observed in all readout measures examined within this study and thus data is presented in aggregate incorporating male and female data.

To determine whether rats fed a HCHF diet became obese, their weights were tracked weekly over the study period from nine to twelve months of age. As expected, the mass of all rats increased over the study period. By the end of the study period, non-transgenic rats (nTg) on CHOW weighed 371 ± 43.2 g while nTg on HCHF diet weighed 464.2 ± 42.8 g (p = 0.005). TgAD on CHOW weighed 437.5 ± 40 g while TgAD on HCHF diet weighed 495.1 ± 36.7 g (p = 0.04).

### Increased myelin and oligodendrocytes in the somatosensory cortex

Previous studies have found that obesity causes myelin damage and a loss of white matter integrity in both males and females of all age groups, with the most pronounced changes in people with body mass indices over thirty^27^. Thus, we investigated whether obesity impacted myelin in nTg and TgAD rats fed either a CHOW or a HCHF diet. We measured myelin density in the striatum and the somatosensory cortex using the myelin-specific Black-Gold II stain. Transgene (p = 0.39), diet (p = 0.59), and the transgene-diet interaction (p = 0.19) had no significant effects on myelin density in the striatum (Fig. 1a,b). However, diet (p = 0.02) and the transgene-diet interaction (p = 0.0002) had significant effects on myelin density in the somatosensory cortex, while transgene did not (p = 0.09) (Fig. 1c,d). nTg rats on HCHF or CHOW had no differences in myelin density (p = 0.77) while TgAD rats on HCHF diet had a 42.5 ± 7.94% increase in myelin density compared to TgAD on CHOW (p < 0.0001) (Fig. 1d). These data indicate that myelin density increases in response to the interaction between HCHF diet and AD transgene in the somatosensory cortex but not the striatum.

**Figure 1:**
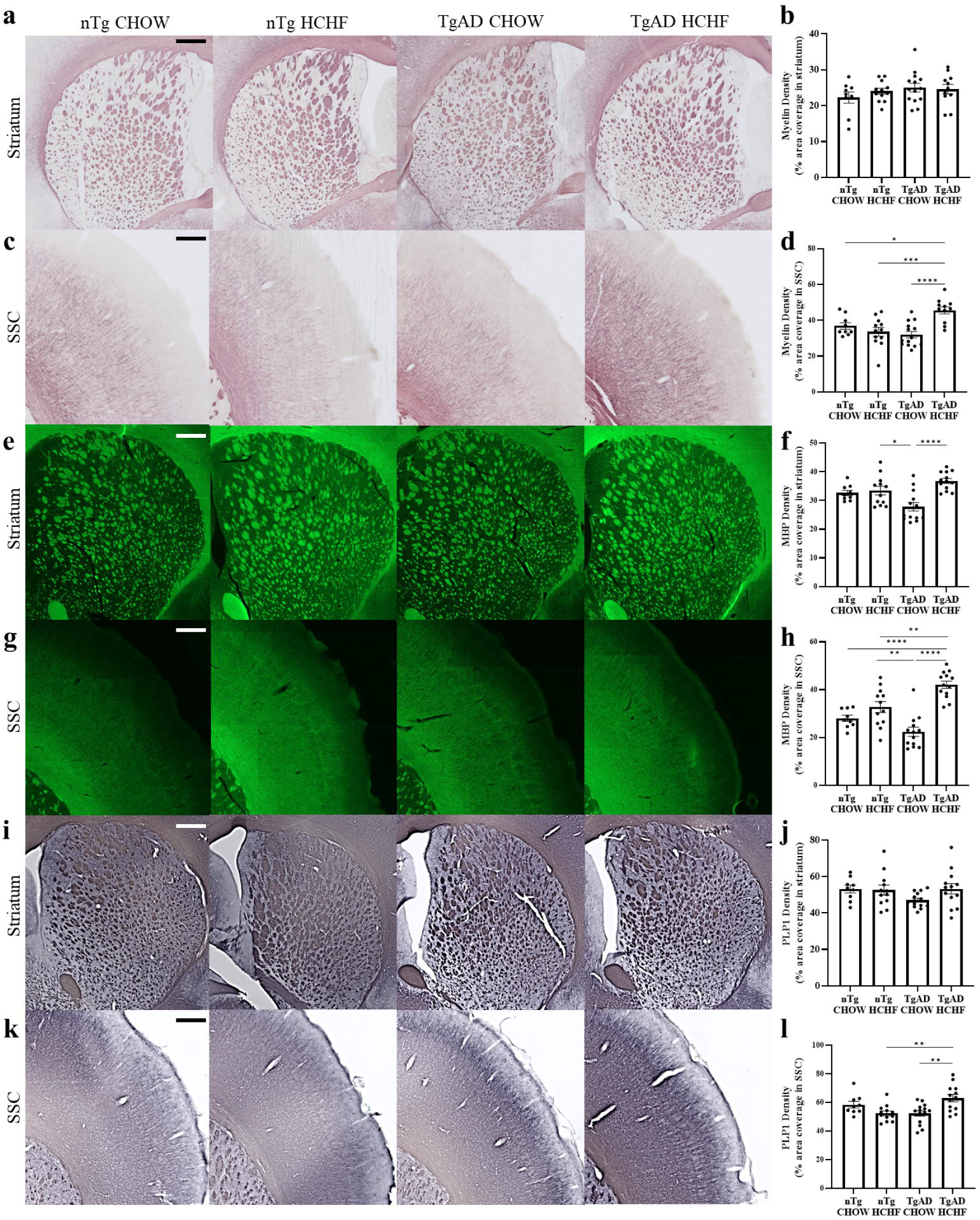
Increased myelin in TgAD HCHF in somatosensory cortex. (**a**) Representative images of Black Gold II-stained myelin in striatum. Scale bar=700 μm. (**b**) Quantification of Black-Gold II stain in striatum suggests no transgene-diet interaction (transgene*diet p=0.39; transgene p=0.19; diet p=0.59). (**c**) Representative images of Black Gold II-stained myelin in somatosensory cortex. Scale bar=375 μm. (**d**) Quantification of Black-Gold II stain in somatosensory cortex suggests a transgene-diet interaction and diet increased myelin density (transgene*diet p=0.002; transgene p=0.09; diet p=0.02; nTg CHOW vs. nTg HCHF p=0.70; TgAD CHOW vs. TgAD HCHF p<0.0001; nTg CHOW vs. TgAD HCHF p=0.04; nTg HCHF vs. TgAD HCHF p=0.0009). (**e**) Representative images of MBP immunoreactivity in striatum. Scale bar=500 µm. (**f**) Quantification of MBP immunoreactivity in striatum suggests a transgene-diet interaction and diet increased MBP immunoreactivity (transgene*diet p=0.03; transgene p=0.58; diet p=0.0004; nTg CHOW vs. nTg HCHF p=0.97; nTg HCHF vs. TgAD CHOW p=0.01; TgAD CHOW vs. TgAD HCHF p<0.0001). (**g**) Representative images of MBP immunoreactivity in somatosensory cortex. Scale bar=350 μm. (**h**) Quantification of MBP immunoreactivity in somatosensory cortex suggests transgene-diet interaction and diet increased MBP density (transgene*diet p=0.0003; transgene p=0.34; diet p<0.0001; nTg CHOW vs. nTg HCHF p=0.35; nTg CHOW vs. TgAD HCHF p<0.0001; nTg HCHF vs. TgAD CHOW p=0.001; nTg HCHF vs. TgAD HCHF p=0.004; TgAD CHOW vs. TgAD HCHF p<0.0001). (**i**) Representative images of PLP1 immunoreactivity in striatum. Scale bar=550 μm. (**j**) PLP1 quantification in striatum shows no impact of diet or transgene (transgene*diet p=0.15; transgene p=0.25; diet p=0.23). (**k**) Representative images of PLP1 immunoreactivity in somatosensory cortex. Scale bar =325 μm. (**l**) Quantification of PLP1 immunoreactivity in somatosensory cortex suggests a transgene-diet interaction increased PLP1 density in TgAD fed HCHF rats (transgene*diet p=0.0004; transgene p=0.31; diet p=0.27; nTg CHOW vs. nTg HCHF p=0.29; TgAD CHOW vs. TgAD HCHF p=0.003; nTg HCHF vs. TgAD HCHF p=0.005). n=9 (nTg CHOW), 12 (nTg HCHF), 14 (TgAD CHOW), and (TgAD HCHF). Two-way ANOVA with *post-hoc* Tukey’s HSD test. * denotes p<0.05, ** denotes p<0.01, *** denotes p<0.001, and **** denotes p<0.0001.

To further investigate the impact of HCHF and obesity on myelin, we measured the immunoreactivity of two myelin proteins, myelin-basic protein (MBP) and myelin proteolipid protein 1 (PLP1) in the striatum and the somatosensory cortex. MBP maintains the structure of myelin and is crucial to the myelination process^28^. Diet (p = 0.0004) and the transgene-diet interaction (p = 0.003) had significant effects on MBP immunoreactivity in the striatum while transgene did not (p = 0.58) (Fig. 1e,f). nTg on HCHF had no differences compared to nTg on CHOW (p = 0.97) (Fig. 1e,f). Surprisingly, despite no change in myelin density in the striatum, TgAD on HCHF had increased MBP immunoreactivity compared to TgAD on CHOW in this region (33.2 ± 4.71%; p < 0.0001) (Fig. 1e,f). In the somatosensory cortex, diet (p < 0.0001) and the transgene-diet interaction (p = 0.0003) had significant impacts on MBP immunoreactivity, while transgene did not (p = 0.34) (Fig. 1g,h). nTg on HCHF had no changes in MBP immunoreactivity compared to nTg on CHOW (p = 0.36) in the somatosensory cortex (Fig. 1g,h). TgAD on HCHF had increased MBP immunoreactivity compared to TgAD on CHOW within the somatosensory cortex (88.1 ± 20.1%; p < 0.0001) (Fig. 1g,h). To verify the increased myelin in the somatosensory cortex, we measured the immunoreactivity of PLP1, which also stabilizes myelin structure and promotes oligodendrocyte development^29^. In agreement with the MBP immunoreactivity, transgene (p = 0.25), diet (p = 0.23), and the transgene-diet interaction (p = 0.15) had no significant impact on PLP1 in the striatum. (Fig. 1i,j). In the somatosensory cortex, the transgene-diet interaction was significant (p = 0.0004), while diet (p = 0.27) and transgene (p = 0.31) were not (Fig. 1k,l). Within the somatosensory cortex, nTg on HCHF diet had no changes in PLP1 immunoreactivity compared to nTg on CHOW (p = 0.29) in the somatosensory cortex (Fig. 1k,l). TgAD on HCHF had increased PLP1 immunoreactivity compared to TgAD on CHOW (20.6 ± 3.01%; p = 0.003) in the somatosensory cortex. Together, these results demonstrate an interaction between transgene and diet that increases myelin density and myelin-related proteins in the somatosensory cortex.

Oligodendrocytes are the myelin-producing cells of the central nervous system and, in AD, participate in defective myelination and undergo apoptosis^30^. Since we observed changes in myelination in the striatum and somatosensory cortex, we next quantified oligodendrocytes using the oligodendrocyte-specific marker, Olig2. Transgene (p = 0.64), diet (p = 0.78), and the transgene-diet interaction (p = 0.17) had no significant impacts on the number of Olig2-positive cells in the striatum (Fig. 2a,b). On the other hand, transgene (p < 0.0001), diet (p = 0.0001), and the transgene-diet interaction (p < 0.0001) were significant factors in Olig2 counts such that TgAD on HCHF diet had significantly more Olig2+ cells compared to TgAD on CHOW in the somatosensory cortex (19.3 ± 1.27%; p < 0.0001) while nTg on HCHF diet had no changes in Olig2 compared to nTg on CHOW (p = 0.85) (Fig. 2c,d). Overall, these results suggest an increase in oligodendrocytes in the somatosensory cortex due to a HCHF diet and AD progression.

**Figure 2:**
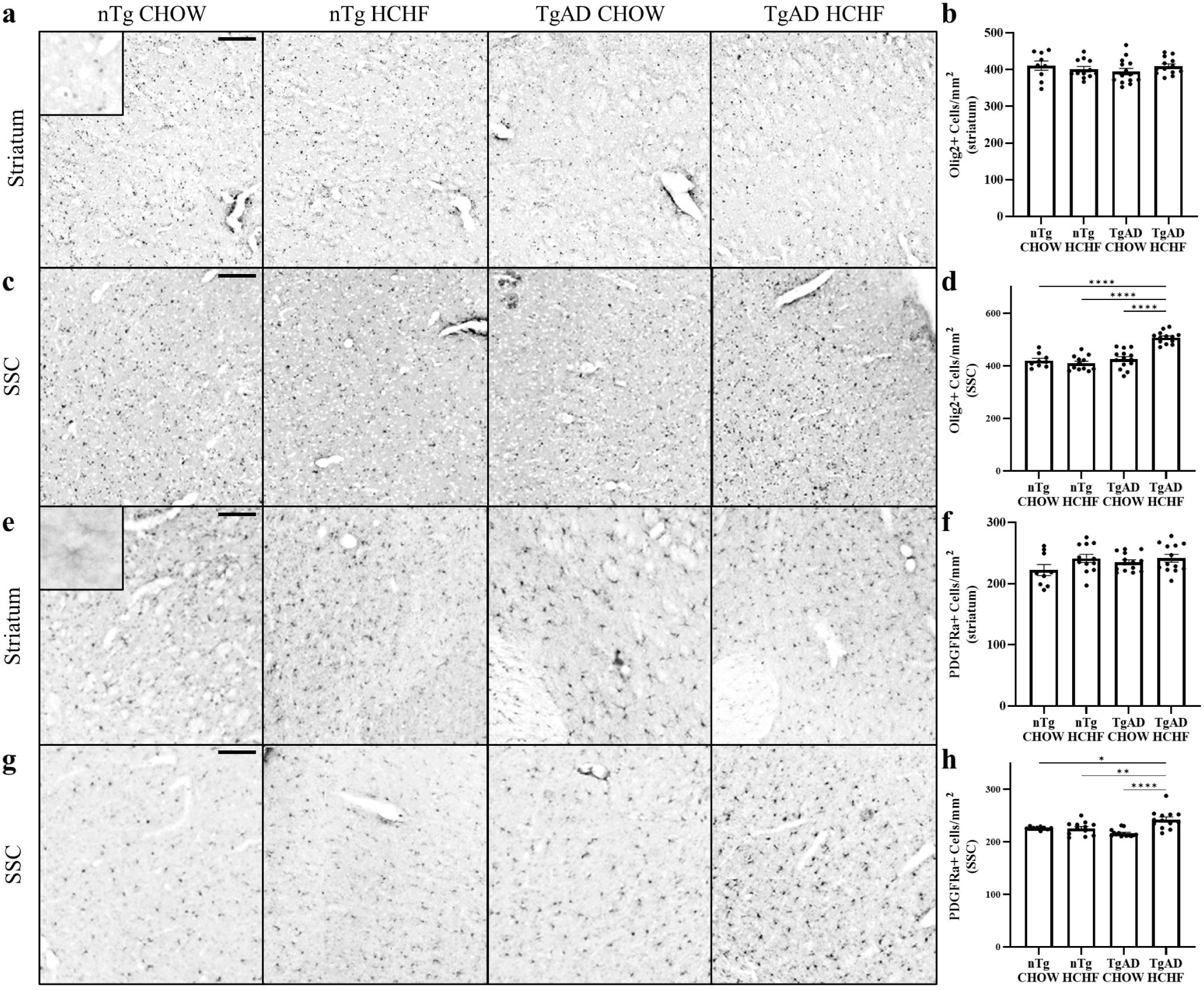
Increased oligodendrocytes and oligodendrocyte progenitor cells in TgAD on HCHF in the somatosensory cortex. (**a**) Representative images of Olig2+ cell immunoreactivity in the striatum. Scale bar = 150 μm. (**b**) Olig2+ cell quantification in the striatum shows no transgene-diet interaction (transgene*diet p=0.17; transgene p=0.64; diet p=0.78; nTg CHOW vs. nTg HCHF p=0.88; TgAD CHOW vs. TgAD HCHF p=0.59). (**c**) Representative images of Olig2+ cell immunoreactivity in the somatosensory cortex. Scale bar = 125 μm. (**d**) Quantification of Olig2+ cell immunoreactivity in the somatosensory cortex suggests a transgene-diet interaction increased oligodendrocyte counts in TgAD fed HCHF rats (transgene*diet p<0.0001; transgene p<0.0001; diet p=0.0001; nTg CHOW vs. nTg HCHF p=0.85; nTg CHOW vs. TgAD HCHF p<0.0001; nTg HCHF vs. TgAD HCHF p<0.0001; TgAD CHOW vs. TgAD HCHF p<0.0001). (**e**) Representative images of PGDGRα+ cell immunoreactivity in the striatum. Scale bar = 150 μm. (**f**) PDGFRα+ cell quantification shows no transgene-diet interaction in the striatum (transgene*diet p=0.33; transgene p=0.31; diet p=0.05; nTg CHOW vs. nTg HCHF p=0.20; TgAD CHOW vs. TgAD HCHF p=0.86). (**g**) Representative images of PGDGRα+ cell immunoreactivity in the somatosensory cortex. Scale bar = 150 μm. (**h**) Quantification of PDGFRα+ cell immunoreactivity in the somatosensory cortex suggests a transgene-diet interaction increased oligodendrocyte progenitor cell counts in TgAD fed HCHF rats (transgene*diet p=0.0005; transgene p=0.32; diet p=0.0008; nTg CHOW vs. nTg HCHF p=0.999; nTg CHOW vs. TgAD HCHF p=0.02; nTg HCHF vs. TgAD HCHF p=0.006; TgAD CHOW vs. TgAD HCHF p<0.0001). n = 9 (nTg CHOW), 12 (nTg HCHF), 14 (TgAD CHOW), and 13 (TgAD HCHF). Two-way ANOVA with *post-hoc* Tukey’s HSD test. * denotes p < 0.05, ** denotes p < 0.01, *** denotes p < 0.001, and **** denotes p < 0.0001.

AD pathology causes oligodendrocyte progenitor cells to undergo apoptosis, which compromises myelin integrity^30^. Since we observed an increase in oligodendrocytes, oligodendrocyte progenitor cells were quantified using the oligodendrocyte progenitor cell-specific marker, PDGFRα. In the striatum, diet (p = 0.04) had a significant effect on the number of PDGFRα+ cells while transgene (p = 0.31) and the transgene-diet interaction (p = 0.33) did not (Fig. 2e,f). However, no significant differences were found between TgAD on HCHF compared to TgAD on CHOW (p = 0.86) or nTg on HCHF compared to nTg on CHOW (p = 0.20) (Fig. 2e,f). In the somatosensory cortex, diet (p = 0.0008) and the transgene-diet interaction (p = 0.0005) significantly impacted the number of PDGFRα+ cells such that TgAD on HCHF diet had significantly more PDGFRα+ cells than TgAD on CHOW (11.97 ± 0.66%; p < 0.0001) (Fig. 2g,h). However, in the somatosensory cortex transgene (p = 0.32) had no significant impact on PDGFRα+ cells and there were no differences between nTg on HCHF compared to nTg on CHOW (p = 0.999) (Fig. 2g,h). Overall, these data demonstrate an increase in oligodendrocytes and oligodendrocyte progenitor cells in the somatosensory cortex that may have led to the increase in myelin density as a result of HFHC diet.

### No neuronal loss, decreased inflammation in the somatosensory cortex

Oligodendrocytes and neurons interact at the axon to direct myelination. When this interaction is interrupted, neurodegeneration may occur^31^. AD is characterized by a loss of neurons^22,23^ and thus we examined whether HCHF diet impacted neuronal loss using neuronal nuclei (NeuN) immunoreactivity. In the striatum, transgene (p = 0.06), diet (p = 0.55), and the transgene-diet interaction (p = 0.27) had no significant effects on NeuN immunoreactivity (Fig. 3a,b). Similarly in the somatosensory cortex, transgene (p = 0.07), diet (p = 0.33), and the transgene-diet interaction (p = 0.25) had no significant effects on NeuN immunoreactivity (Fig. 3c,d) This suggests that the HCHF diet does not contribute to neuronal loss in the striatum and the somatosensory cortex.

**Figure 3:**
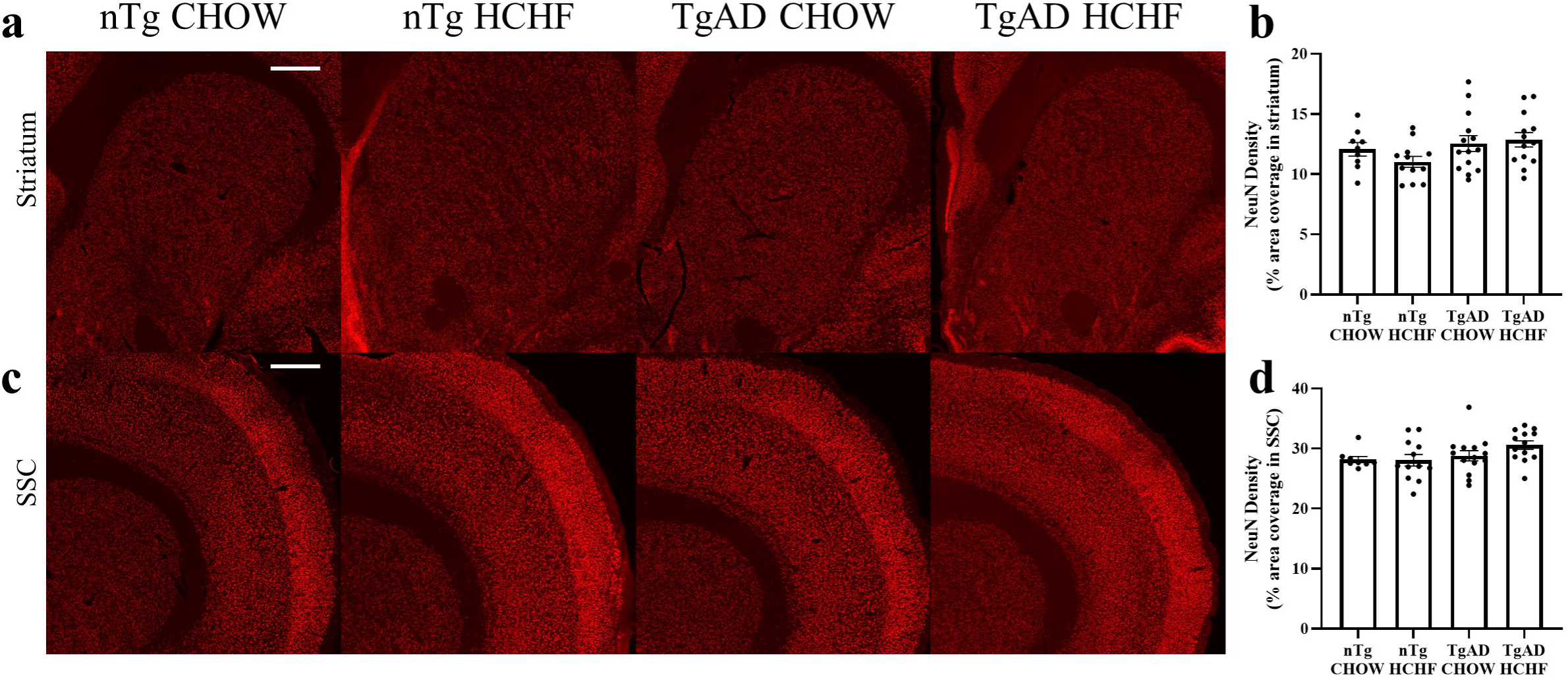
No changes in NeuN expression. (**a**) Representative images show NeuN immunoreactivity in the striatum. Scale bar = 625 μm. (**b**) NeuN immunoreactivity quantification in the striatum shows no significant transgene-diet interaction effects on NeuN expression (transgene*diet p=0.27; transgene p=0.06; diet p=0.55; nTg CHOW vs. nTg HCHF p=0.66; TgAD CHOW vs. TgAD HCHF p=0.98). (**c**) Representative images show NeuN immunoreactivity in the somatosensory cortex. Scale bar = 625 μm. (**d**) NeuN immunoreactivity quantification in the somatosensory cortex shows no significant transgene-diet interaction effects on NeuN expression (transgene*diet p=0.25; transgene p=0.07; diet p=0.33; nTg CHOW vs. nTg HCHF p=0.99; TgAD CHOW vs. TgAD HCHF p=0.37). n = 9 (nTg on CHOW), 12 (nTg on HCHF), 14 (TgAD on CHOW), and 13 (TgAD on HCHF). Two-way ANOVA with *post-hoc* Tukey’s HSD test.

Microglia ensure proper oligodendrocyte and oligodendrocyte progenitor cell development, while contributing to normal myelination^32^. In contrast, activated microglia are a hallmark of AD progression and contribute to neuroinflammation, neuronal loss, and cognitive deficits^33^. Previously, obesity has been found to increase microglial activation^20^. We set out to determine whether HCHF diet and AD progression impacted microglia by measuring ionized calcium binding adaptor molecule 1 (Iba1) immunoreactivity. Increased Iba1 immunoreactivity results from an increase in activated microglia^34^. In the striatum, transgene (p = 0.003) significantly increased Iba1 immunoreactivity while diet (p = 0.005) significantly reduced Iba1 immunoreactivity (Fig. 4a,b). However, there was no transgene-diet interaction (p = 0.97) and so TgAD on HCHF compared to TgAD on CHOW (p = 0.14) and nTg on HCHF compared to nTg on CHOW (p = 0.21) had no significant changes in Iba1 immunoreactivity in the striatum (Fig. 4a,b). In the somatosensory cortex, transgene (p = 0.0006) significantly increased Iba1 immunoreactivity, diet (p < 0.0001) significantly reduced Iba1 immunoreactivity, and the transgene-diet interaction (p = 0.002) significantly impacted Iba1 immunoreactivity such that TgAD on HCHF diet had 24.0 ± 2.73% less Iba1 immunoreactivity compared to TgAD on CHOW in the somatosensory cortex (p < 0.0001) (Fig. 4c,d). However, nTg on HCHF had no significant changes in Iba1 immunoreactivity compared to nTg on CHOW (p = 0.82) in the somatosensory cortex (Fig. 4c,d). This suggests that the HCHF diet in an AD transgene specific manner decrease Iba1 immunoreactivity in the somatosensory cortex.

**Figure 4:**
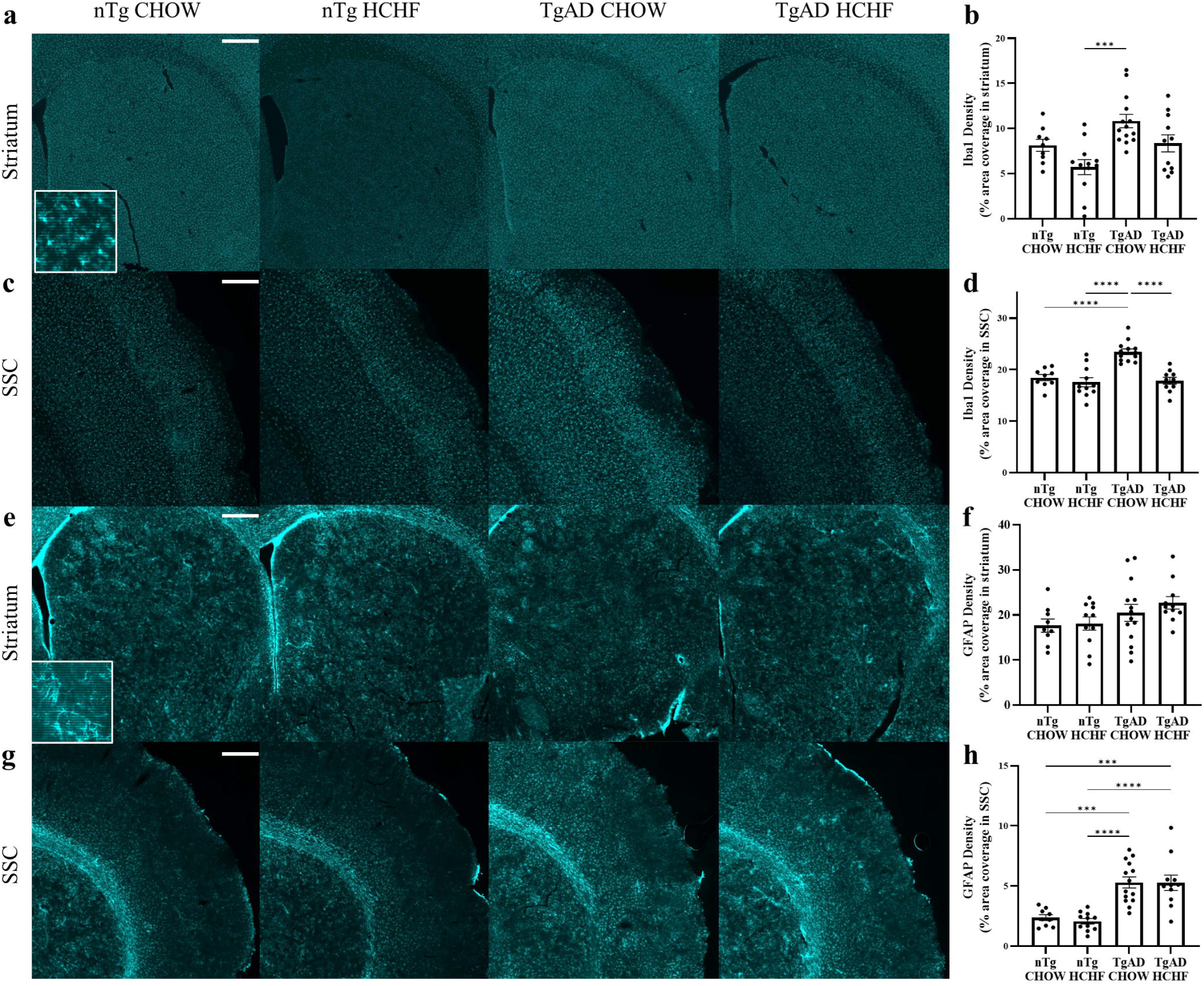
Decreased inflammation in the somatosensory cortex. (**a**) Representative images show Iba1 immunoreactivity in the striatum. Scale bar = 600 μm. (**b**) Iba1 immunoreactivity quantification in the striatum shows no significant transgene-diet interaction (transgene*diet p=0.97; transgene p=0.002; diet p=0.005; nTg CHOW vs. nTg HCHF p=0.21; nTg HCHF vs. TgAD CHOW p=0.0002; TgAD CHOW vs. TgAD HCHF p=0.12). (**c**) Representative images show Iba1 immunoreactivity in the somatosensory cortex. Scale bar = 350 μm. (**d**) Iba1 immunoreactivity quantification in the somatosensory cortex shows a transgene-diet interaction that reduces activated microglia in TgAD HCHF (transgene*diet p=0.002; transgene p=0.0006; diet p<0.0001; nTg CHOW vs. nTg HCHF p=0.81; nTg CHOW vs. TgAD CHOW p<0.0001; nTg HCHF vs. TgAD HCHF p<0.0001; TgAD CHOW vs. TgAD HCHF p<0.0001). n = 9 (nTg on CHOW), 11 (nTg on HCHF), 13 (TgAD on CHOW), and 10 (TgAD on HCHF). Two-way ANOVA with *post-hoc* Tukey’s HSD test. (**e**) Representative images show GFAP immunoreactivity in the striatum. Scale bar = 500 μm. (**f**) GFAP immunoreactivity quantification shows no significant transgene-diet interaction (transgene*diet p=0.60; transgene p=0.03; diet p=0.43; nTg CHOW vs. nTg HCHF p=0.99; TgAD CHOW vs. TgAD HCHF p=0.75). (**g**) Representative images show GFAP immunoreactivity in the somatosensory cortex. Scale bar = 500 μm. (**h**) GFAP immunoreactivity quantification shows no significant transgene-diet interaction (transgene*diet p=0.76; transgene p<0.0001; diet p=0.15; nTg CHOW vs. nTg HCHF p=0.97; nTg CHOW vs. TgAD CHOW p=0.0003; nTg CHOW vs. TgAD HCHF p=0.0006; TgAD CHOW vs. TgAD HCHF p=0.99). n = 9 (nTg on CHOW), 11 (nTg on HCHF), 14 (TgAD on CHOW), and 10 (TgAD on HCHF). Two-way ANOVA with *post-hoc* Tukey’s HSD test. * denotes p < 0.05, ** denotes p < 0.01, *** denotes p < 0.001, and **** denotes p < 0.0001.

Astrocytes and oligodendrocytes interact to induce remyelination^35^. In AD, reactive astrocytes contribute to neuroinflammation and neurodegeneration^36^. We investigated the HCHF diet and AD progression on reactive astrocytes by measuring glial fibrillary acidic protein (GFAP) immunoreactivity. Increased GFAP immunoreactivity is a marker of reactive astrocytes^37^. In the striatum, transgene (p = 0.03) significantly increased GFAP immunoreactivity while diet (p = 0.43) and the transgene-diet interaction had no significant effect on GFAP immunoreactivity (p = 0.60) (Fig. 4e,f). In the striatum, TgAD on HCHF compared to TgAD on CHOW (p = 0.75) and nTg on HCHF compared to nTg on CHOW (p = 0.99) had no significant changes in GFAP immunoreactivity (Fig. 4e,f). Similarly, in the somatosensory cortex, transgene (p < 0.0001) significantly increased GFAP immunoreactivity while diet (p = 0.77) and the transgene-diet interaction (p = 0.76) had no significant effect (Fig. 4g,h). In the somatosensory cortex, TgAD on HCHF compared to TgAD on CHOW (p = 0.99) and nTg on HCHF compared to nTg on CHOW (p = 0.97) had no significant changes in GFAP immunoreactivity (Fig. 4g,h). Together, this suggests that HCHF diet has no effect on astrocyte activation as measured by GFAP immunoreactivity in either the somatosensory cortex or striatum.

### Increased AD pathology in the somatosensory cortex

A key pathological hallmark of AD is the presence of A,B plaques in the brain^21,22^. A,B deposition is directly and indirectly linked to myelin dysfunction in AD, and a high-fat diet has been found to increase A,B plaques^17,38^. Therefore, after observing increased myelin, we examined whether a HCHF diet has an impact on A,B plaques utilizing the Ab-specific antibody, 6F3D. In the striatum, no significant difference was observed in A,B plaques between TgAD on CHOW and TgAD on HCHF (p = 0.91) (Fig. 5a,c). A,B plaques increased by 46.4 ± 19.2% in the somatosensory cortex in TgAD on HCHF compared to TgAD on CHOW (p = 0.04) (Fig. 5b,d). Cerebral amyloid angiopathy which involves A,B deposition in the vasculature is present in up to 90% of AD patients ^39^. Cerebral amyloid angiopathy impairs the brain’s vasculature leading to brain atrophy, cognitive impairment, and ischemia^39^. By compromising the vasculature, cerebral amyloid angiopathy has been found to cause lesions in white matter^40^. We have previously found cerebral amyloid angiopathy in the penetrating arterioles and amyloid accumulation in the venules of the somatosensory cortex^22^. Thus, we set out to determine if cerebral amyloid angiopathy was impacted by diet. TgAD on HCHF had a 10.6 ± 1.28% increase in cerebral amyloid angiopathy in the somatosensory cortex compared to TgAD on CHOW (p = 0.04) (Fig. 5e,f). A,B plaques and cerebral amyloid angiopathy are not detected in nTg rats and thus not quantified herein (data not shown). Together, this shows increased A,B in the somatosensory cortex parenchyma and vasculature due to a HCHF diet.

**Figure 5:**
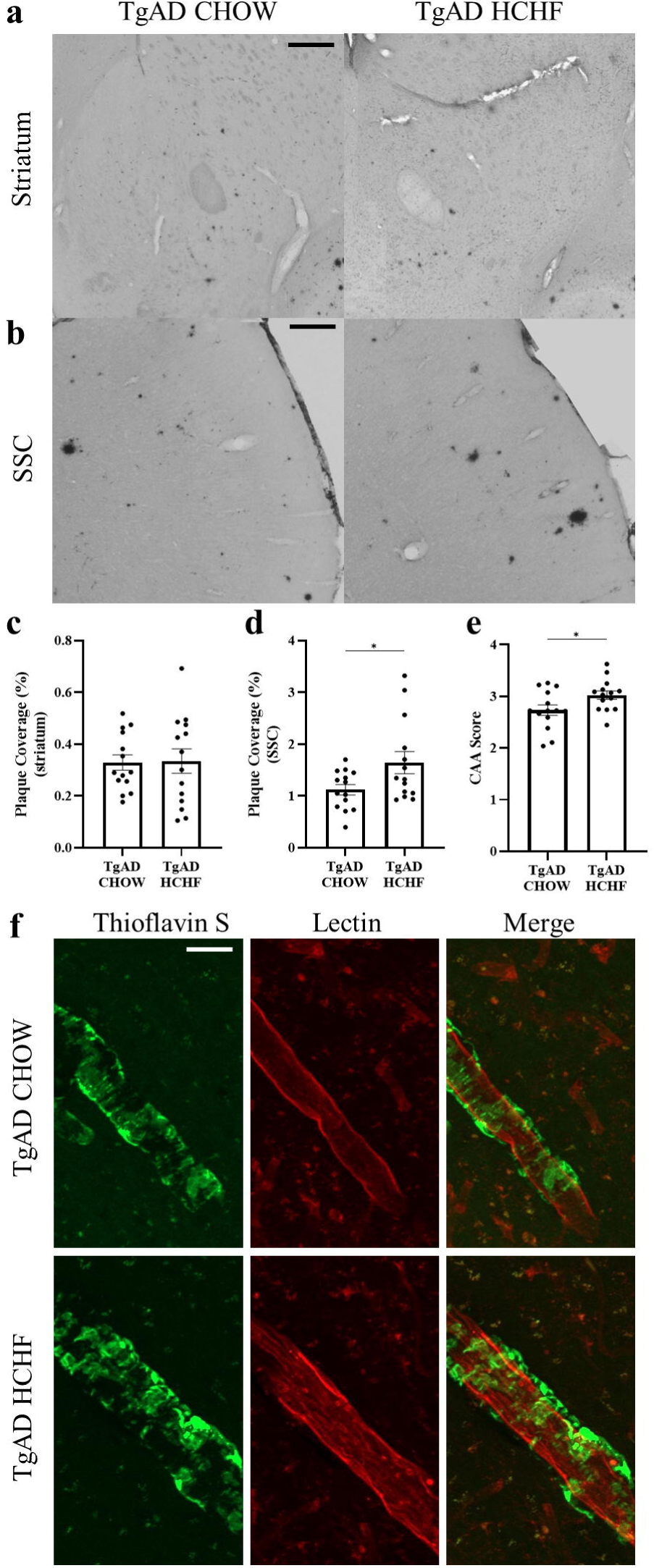
Increased A*β*) in the somatosensory cortex of TgAD on HCHF compared to TgAD on CHOW. (**a**) Representative images show plaque coverage from 6F3D immunoreactivity in the striatum. Scale bar = 425 μm. (**b**) Representative images show 6F3D immunoreactivity in the somatosensory cortex. Scale bar = 300 μm. (**c**) 6F3D immunoreactivity quantification in the striatum shows no significant effect of HCHF on A,B plaque load (p=0.91). (**d**) 6F3D immunoreactivity quantification in the somatosensory cortex suggests that HCHF increases A,B plaque load (p=0.04). (**e**) Quantification of cerebral amyloid angiopathy (CAA) suggests that HCHF increases vascular amyloid deposition in TgAD rats (p=0.04). (**f**) Representative images of a thioflavin/lectin fluorescent co-stain show CAA in the somatosensory cortex. Scale bar = 25 μm. n = 14 (TgAD on CHOW), and 13 (TgAD on HCHF) with an unpaired *t*-test. * denotes p < 0.05.

Intracellular tau inclusions contribute to neurodegeneration in AD and are frequently colocalized with A,B plaques^41^. Thus, we set out to determine if tau inclusions increase in conjunction with the increase in A,B plaques. We examined tau inclusions using the disease-specific antibody, paired helical filament-1 (PHF-1) immunoreactivity. We analyzed both plaque-associated and non-plaque associated tau inclusions (Fig 6). In the striatum, plaque-associated (p = 0.33) and non-plaque associated inclusions (p = 0.49) remained unchanged between TgAD on CHOW and TgAD on HCHF diets (Fig. 6a,b). In the somatosensory cortex, there was a 48.5 ± 9.55% increase in the number of plaque-associated tau inclusions of TgAD on HCHF compared to TgAD on CHOW (p < 0.0001), with no significant differences in non-plaque associated tau inclusions (p = 0.30) (Fig. 6c,d). Overall, this shows that a HCHF diet increases AD pathology in the somatosensory cortex.

**Figure 6:**
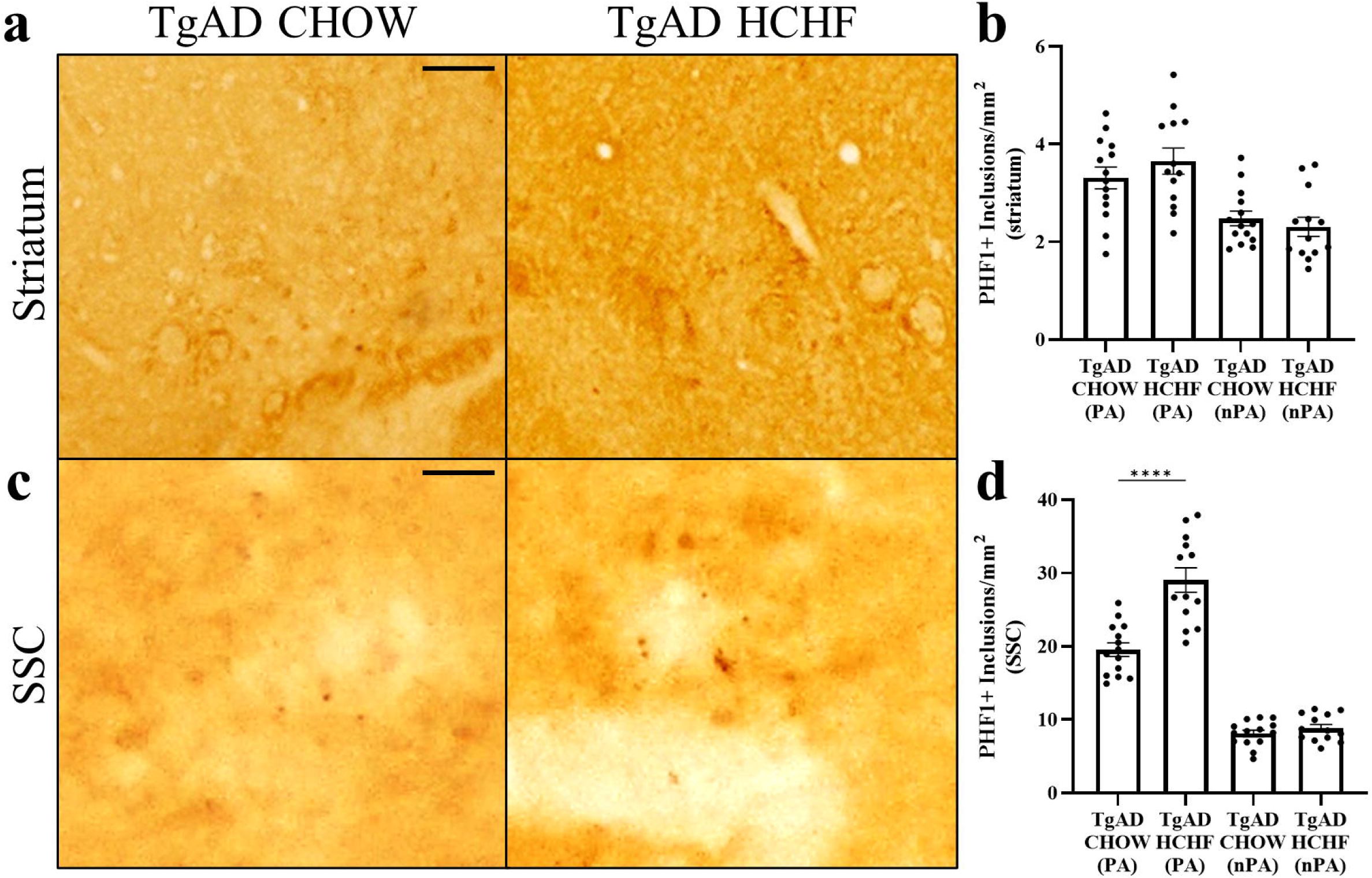
Increased plaque-associated tau in the somatosensory cortex of TgAD on HCHF. (**a**) Representative images of a PHF-1 stain shows tau inclusions in the striatum. Scale bar = 100 μm. (**b**) Quantification of PHF-1 shows no changes in plaque-associated (PA) and non-plaque associated (nPA) tau in the striatum (p=0.33; p=0.49). (**c**) Representative images of a PHF-1 stain show tau inclusions in the somatosensory cortex. Scale bar = 20 μm. (**d**) PHF-1 quantification suggests that HCHF increases the number of plaque-associated tau in the somatosensory cortex (p<0.0001), but not non-plaque associated tau (p=0.30). n = 14 (TgAD on CHOW), and 13 (TgAD on HCHF) with an unpaired *t*-test. **** denotes p < 0.0001.

## 4 Discussion

Herein, we showed that a varied HCHF diet induced obesity in both TgAD and nTg rats from nine to twelve months of age. A HCHF diet in TgAD led to larger effect size outcomes in the somatosensory cortex compared to the striatum, confirming region-specific effects of a HCHF diet. We show that HCHF diet-induced obesity increases myelin within the somatosensory cortex with a corresponding increase in oligodendrocytes and oligodendrocyte progenitor cells. We observed no loss in neurons. Further, there was decreased activated microglia, yet an increase in A,B, cerebral amyloid angiopathy, and plaque-associated tau in the somatosensory cortex. These combined results suggests that diet effects on AD pathology are independent from other beneficial effects on myelin in the somatosensory cortex^42–44^.

Restoring myelin loss may be a promising method of reversing cognitive decline in AD^45^. Several studies have reported myelin loss in AD as a result of A,B toxicity and tauopathy^46,47^. The loss in myelin exacerbated cognitive decline in human^48^. Further during normal aging in humans, myelin loss has been associated with age-related cognitive decline^49,50^. The increase in myelin that we report in the obese TgAD rats may provide one mechanism leading to the obesity paradox observed in aged people. An increase in myelin can rescue cognitive deficits in AD rodent models^44^. Chen et al. utilized genetic knockouts and antagonists of the muscarinic receptor 1, both of which promoted oligodendrocyte progenitor cell differentiation^44^. Oligodendrocyte progenitor cell differentiation led to an increase in oligodendrocytes, which enhanced myelin renewal in mice^44^. Increased myelin was found to rescue memory and improved neuronal function^44^. Increased MBP in both striatum and somatosensory cortex may also contribute to these benefits. AD patients have decreased MBP, which is consistent with the decreased levels of myelin^51^. Thus, by increasing and stabilizing myelin expression, neuronal function and cognitive deficits in AD may be attenuated. Notably, oligodendrocytes require significant amounts of energy to increase myelination^52^. One possible source of this energy is the HCHF diet consumed by the rats in our study, which is abundant in energy^53^. Another benefit of a HCHF diet is the diverse carbohydrate composition. Increasing carbohydrates, such as glucose and lactate, has been shown to increase oligodendrocyte development and myelination in rats^54^. Since oligodendrocyte progenitor cells continue to divide and proliferate throughout adulthood, the observed increase in PDGFRα+ cells may also contribute to the increase in oligodendrocytes^55^. Overall, our results suggest that a HCHF diet increases the number of oligodendrocytes, which increase myelination.

Our overall results show that a HCHF diet has a larger effect size on the somatosensory cortex compared to the striatum. This suggests that regions with less white matter may be more positively affected by a HCHF diet compared to regions dense in white matter. In human AD, both white and gray matter are impacted, however, damage to regions dominated by gray matter tend to be more severe^56,57^. One possible explanation for the observed increase in myelin in the somatosensory cortex is that gray matter contains axons that are scarcely myelinated. In contrast, white matter is heavily myelinated leaving little space for more myelin. This extra space in the gray matter may accommodate an increase in myelin. Functionally, the striatum acts as a relay between the thalamus and the cerebral cortex, thus, the dense myelin of the striatum is important for efficient communication.^58^. However, the somatosensory cortex is involved in cognitive processes like motor and haptic memory^26^. Since myelination is important to preserve memories in the long-term, increased myelin may be the result of improved memory consolidation^59^. These functional differences may help to explain the larger effect size within the somatosensory cortex due to a HCHF diet compared to the striatum.

A limitation of this study is the lack of behavioral data for the somatosensory cortex and striatum. Although we observed beneficial changes in the somatosensory cortex at the protein level, it remains unclear if increased myelination in the somatosensory cortex has effects at the behavioral level.

However, our results do suggest that the HCHF diet and the increased myelin are not harmful to the somatosensory cortex. We observe no changes in NeuN expression and decreased microglial activation, which indicates that there was no additional damage to brain tissue.

In conclusion, we show that obesity through a HCHF diet in early stages of AD progression has beneficial outcomes with larger effect sizes in the somatosensory cortex than the striatum. Due to the diverse range of foods used in the HCHF diet, we are simultaneously targeting several metabolic pathways^53^. By identifying and targeting these pathways, we can stimulate myelination in AD. Further, our results suggest that the obesity paradox may in part be due to changes in myelin. Obesity during early stages of AD progression increases myelin which may prevent/slow cognitive decline and reduce the risk of AD. As a result, we present one mechanism that may contribute to the obesity paradox.

## 5 Methods

### Non-transgenic and Transgenic Rats

nTg and TgAD rats were outbred on a Fischer strain and housed on a 12-hour light:12-hour dark schedule with *ad libitum* access to food and water. TgAD rats overexpress human amyloid precursor protein containing the Swedish mutation and an exon 9 deletion of Presenilin 1, both driven by a mouse prion promoter^21^. Ethical approval of all experimental procedures was granted by The Animal Care Committee of the Sunnybrook Health Sciences Center, which adheres to the Policies and Guidelines of the Canadian Council on Animal Care, the Animals for Research Act of the Provincial Statute of Ontario, the Federal Health of Animals Act, and is compliant with ARRIVE guidelines.

### Diet

Rats fed CHOW were given standard chow and water ad libitum. Rats fed a HCHF diet were given the choice between standard chow and three HCHF food items *ad libitum* starting at 9 months of age. The HCHF rats were also given the choice between water and 12% sucrose solution to mimic soda consumption. The three HCHF items were rotated with three different HCHF items three times a week to maintain appetite and palatability. Food intake was tracked by weighing leftover food after the three HCHF items were rotated. Rats were weighed weekly.

### Tissue collection

The nTg and TgAD rats were sacrificed under 5% isoflurane at 12 months of age. The rats were transcardially perfused with PBS-0.1% heparin followed by PBS-4% paraformaldehyde. Brains were extracted and post-fixed overnight in PBS-4% paraformaldehyde. The brains were cryopreserved in PBS-30% sucrose. Tissue was collected using a sliding microtome and 40 µm sections were collected from bregma +4.00 to -6.00. Four evenly spaced sections from bregma +2.00 to -1.00 were used for all pathological readout measures.

### Immunofluorescence for NeuN, MBP, Iba1, GFAP, Lectin, and Thioflavin

All washes were in 1X PBS and occurred between all antibody incubations. Sections were blocked in 0.5% Triton X-100 and 0.5% bovine serum albumin in PBS for 1 hour at room temperature. Sections were incubated with primary antibody diluted in block overnight at 4L. The sections were incubated with secondary antibody diluted in block. Sections were mounted and coverslipped using PVA DABCO (Sigma-Aldrich #10981). The following primary antibodies were used: guinea pig anti-NeuN (Millipore #ABN90; 1:500), rat anti-MBP (Millipore #MAB386; 1:50), rabbit anti-Iba1 (WAKO #019-19741; 1:500), fluorescent-tagged tomato lectin (Vector #DL1177; 1:200), and mouse anti-GFAP (abcam #ab190288; 1:1000). The following secondary antibodies were used: anti-guinea pig (Invitrogen #A11076; 1:250), anti-rat (Invitrogen #A11006; 1:250), anti-rabbit (Invitrogen #A31573; 1:250), and anti-mouse (Invitrogen #A21050; 1:250).

For the thioflavin-S stain, sections were washed in 1X PBS. Sections were incubated in 1% Thioflavin-S (Sigma-Aldrich #T1892) in distilled water for 8 minutes at room temperature. Sections were washed twice in 70% ethanol followed by three washes in 1X PBS. Sections were then processed for further immunofluorescent co-staining.

### Immunohistochemistry for PLP1, Olig2, PDGFRα, Aβ, and Tau

Olig2: All washes were in 1X PBS and were done between all antibody incubations, unless otherwise stated. Sections were incubated in 3% hydrogen peroxide in PBS for 30 minutes at room temperature. The blocking step and antibody dilutions were in 0.5% Triton X-100 and 0.5% BSA in PBS. The sections were blocked for 1 hour at room temperature and then without washing, incubated with rabbit anti-Olig2 (abcam #ab109186; 1:200) overnight at room temperature. The sections were incubated with Vectastain ABC kit anti-rabbit secondary antibody (Vector Laboratories #PK-4001; 1:400) for 1.5 hours at room temperature. After washing, the sections were incubated with ABC solution (Vector Laboratories #PK-4001; 1:400) diluted in PBS for 1 hour at room temperature.

Sections were developed using 3,3’-diaminobenzidine peroxidase substrate kit (Vector Laboratories #SK-4100). The sections were mounted, dehydrated in an ascending ethanol series (70%, 95%, 100%) and xylene, and then cover slipped using Cytoseal.

PLP1 and PDGFRα: The protocol is similar to Olig2, however, there was an additional antigen retrieval step prior to blocking. The sections underwent antigen retrieval incubated in 10 mM sodium citrate, pH 6.0, for 45 minutes at 85°C. ABC solution was diluted at 1:200. Primary antibodies used were mouse anti-PLP1 (Biorad #MCA839G; 1:250) and goat anti-PDGFRα (R&D Systems #AF1062; 1:100). Secondary antibodies used were Vectastain ABC kit anti-mouse secondary antibody (Vector Laboratories #PK-4002; 1:400) and Biotin-SP AffiniPure Donkey Anti-Goat IgG (Jackson Laboratories #705-065-147; 1:1000).

Aβ: The protocol is similar to Olig2, however, there was an additional antigen retrieval step prior to blocking. The sections underwent antigen retrieval incubated in 70% formic acid for 5 minutes at room temperature. ABC solution was diluted at 1:200. The primary antibody was mouse anti-human Aβ (Dako #M0872; 1:400) and the secondary antibody was Vectastain ABC kit anti-mouse secondary antibody (Vector Laboratories #PK-4002; 1:400).

Tau: All washes were in tris-buffered saline (TBS) and were conducted between all antibody incubations. 0.05% Triton X-100 was added to TBS following primary incubation until 3,3’-diaminobenzidine development. Sections were incubated in 3% hydrogen peroxide in 0.25% Triton X-100 TBS for 30 minutes at room temperature. The sections were blocked in 5% skim milk in TBS for 1 hour at room temperature and without washing, incubated with mouse IgG1 anti-PHF1 (a gift from Dr. Peter Davies; 1:1000) in 5% skim milk in TBS overnight at 4L. The sections were incubated with goat anti-mouse IgG1-biotinXX antibody (Invitrogen #A10519; 1:200) in 20% superblock in TBS for 1.5 hours at room temperature. The sections were incubated with ABC solution (Vector Laboratories #PK-4002; 1:200) diluted in 20% superblock in TBS for 1 hour at room temperature. Sections were developed using 3,3’-diaminobenzidine peroxidase substrate kit (Vector Laboratories #SK-4100). The sections were mounted, dehydrated in an ascending ethanol series (70%, 95%, 100%) and xylene, and then cover slipped using Cytoseal.

### Myelin Staining

Myelin fibers were stained using Black Gold II (HistoChem #2BGII). Sections were washed in saline and then mounted on slides. After rehydrating in saline, the slides were incubated with 1 mL of 0.3% Black Gold II in saline on a heating block at 60L for 6 minutes. The slides were washed in distilled water and fixed in 1% sodium thiosulfate in saline. The slides were washed in distilled water then dehydrated in an ascending alcohol series (70%, 95%, 100%) and xylene, and then cover slipped with Cytoseal.

### Imaging and Statistical Analyses

Whole brain images were taken using a Zeiss Observer.Z1. ImageJ was used to quantify the density of myelin and fluorescent labeling. Regions of interest were outlined to include either the somatosensory cortex or the striatum. These images were set to a background of 12-14 pixels per 1000 µm^2^. Area coverage was measured as a % area. To count the number of Olig2+ cells, objects between 25 pixels^2^ and 300 pixels^2^ were summed after thresholding. To count the number of PDGFRα+ cells, an Ilastik program was trained to recognize cells and images were exported as pixel probabilities. After thresholding the pixel probabilities, objects greater than 25 pixels^2^ were summed. To measure cerebral amyloid angiopathy, a semi-quantitative cerebral amyloid angiopathy score was used as previously established^60^. In brief, a score of 1 was given to blood vessels with 0-20% thioflavin-S coverage, a score of 2 was given to vessels with 21-40% thioflavin-S coverage, up to a maximum score of 5. Tau was quantified by a manual count on ImageJ after adjusting brightness and contrast. Examiner was blind to sex, diet, and transgene throughout the analysis. Using Grubb’s test, one TgAD HCHF rat was a significant outlier across several assays and was removed from the analysis. For activated microglia (1 nTg HCHF, 1 TgAD CHOW, 3 TgAD HCHF) and reactive astrocyte (1 nTg HCHF, 3 TgAD HCHF) analysis, some rats were removed from analysis due to a poor perfusion. GraphPad Prism 9.5.0 was used to perform all statistical analysis and to generate graphs. Two-way ANOVA with a *post-hoc* Tukey’s HSD test was used for all analyses where transgene and diet were independent variables. Unpaired *t*-test was used where only diet was an independent variable.

## 6 Data Availability

All data generated or analysed during this study are included in this published article.

## 8 Acknowledgments

We thank Tina L. Beckett for technical assistance.

## 9 Author Contributions

MA performed tissue processing experiments and drafted the manuscript. AL and JM designed the study. MH breed the rat colonies, fed the rats, weighed the rats, and prepared brains for processing. AL and JR assisted in tissue processing and provided technical assistance. AA created the PDGFRα Ilastik protocol and assisted with PDGFRα analysis. AL and JM edited the manuscript. All authors contributed to the article and approved the submitted version.

